# Multiple Introductions of the *Mycobacterium tuberculosis* Lineage 2–Beijing into Africa over centuries

**DOI:** 10.1101/413039

**Authors:** Liliana K. Rutaihwa, Fabrizio Menardo, David Stucki, Sebastian M. Gygli, Serej D Ley, Bijaya Malla, Julia Feldmann, Sonia Borrell, Christian Beisel, Kerren Middelkoop, E Jane Carter, Lameck Diero, Marie Ballif, Levan Jugheli, Klaus Reither, Lukas Fenner, Daniela Brites, Sebastien Gagneux

## Abstract

The Lineage 2–Beijing (L2–Beijing) sub-lineage of *Mycobacterium tuberculosis* has received much attention due to its high virulence, fast disease progression, and association with antibiotic resistance. Despite several reports of the recent emergence of L2–Beijing in Africa, no study has investigated the evolutionary history of this sub-lineage on the continent. In this study, we used whole genome sequences of 817 L2 clinical strains from 14 geographical regions globally distributed to investigate the origins and onward spread of this lineage in Africa. Our results reveal multiple introductions of L2–Beijing into Africa linked to independent bacterial populations from East-and Southeast Asia. Bayesian analyses further indicate that these introductions occurred during the past 300 years, with most of these events pre-dating the antibiotic era. Hence, the success of L2–Beijing in Africa is most likely due to its hypervirulence and high transmissibility rather than drug resistance.

## 1 Introduction

Tuberculosis (TB) is mainly caused by a group of closely related bacteria referred to as the *Mycobacterium tuberculosis* Complex (MTBC). The MTBC comprises seven phylogenetic lineages adapted to humans and several lineages adapted to different wild and domestic animal species (Gagneux, 2018). The human-adapted lineages of the MTBC show a distinct geographic distribution, with some “generalist” lineages such as Lineage (L)2 and L4 occurring all around the world and others being geographically restricted “specialist” that include L5, L6 and L7 (Coscolla and Gagneux, 2014; Stucki *et al.*, 2016). Africa is the only continent which is home to all seven human-adapted lineages, including the three “specialist” lineages exclusively found on the continent. Current evidence suggests that the MTBC overall originated in Africa (Gagneux, 2018) and subsequently spread around the globe following human migratory events (Wirth *et al.*, 2008; Comas *et al.*, 2013). The broad distribution of some of the “generalist” lineages and their presence in Africa has been attributed to past exploration, trade and conquest. For instance, an important part of the TB epidemics in sub-Saharan Africa is driven by the generalist Latin–American–Mediterranean (LAM) sublineage of L4, which is postulated to have been introduced to the continent post-European contact (Stucki *et al.*, 2016).

Among the different human-adapted MTBC lineages, the L2–Beijing sublineage has been of particular interest (Merker *et al.*, 2015). L2–Beijing has expanded and emerged worldwide from East Asia; its most likely geographical origin (Luo *et al.*, 2015; Merker *et al.*, 2015). In some parts of the world, the recent emergence of L2–Beijing has been linked to increased transmission (Yang *et al.*, 2012; Holt *et al.*, 2018), high prevalence of multidrug–resistant TB (MDR–TB) (Borrell and Gagneux, 2009), and to social and political instability, resulting into displacement of people and poor health systems (Eldholm *et al.*, 2016). Increasingly, L2–Beijing is also being reported in Africa (Bifani *et al.*, 2002; Affolabi *et al.*, 2009; Gehre *et al.*, 2016; Mbugi *et al.*, 2016), and evidence suggests that L2–Beijing in African regions is becoming more prevalent over time (Cowley *et al.*, 2008; van der Spuy *et al.*, 2009; Glynn *et al.*, 2010). Some authors have hypothesized that the introduction of L2–Beijing into South Africa resulted from the importation of slaves from Southeast Asia during the 17^th^ and 18^th^ centuries and/or the Chinese labor forces arriving in the 1900s (van Helden *et al.*, 2002). Alternatively, in West Africa, the presence of L2–Beijing was proposed to reflect more recent immigration from Asia (Affolabi *et al.*, 2009; Gehre *et al.*, 2016). To a certain extent, the recent expansion of L2–Beijing in parts of Africa has been associated with drug resistance (Githui *et al.*, 2004; Klopper *et al.*, 2013) and higher transmissibility (Guerra-Assunção *et al.*, 2015). In addition, a study in the Gambia showed a faster progression from latent infection to active TB disease in patient house-hold contacts exposed to L2–Beijing (de Jong *et al.*, 2008).

Whilst L2–Beijing seems to be expanding in several regions of Africa, no study has formally investigated the evolutionary history of L2–Beijing on the continent. In this study, we used whole genome sequencing data from a global collection of L2 clinical strains to determine the most likely geographical origin of L2–Beijing in Africa and its spread across the continent.

## 2 Results

### 2.1 Phylogenetic inference of L2 strains

We analyzed a total of 817 L2 genomes originating from 14 geographical regions including Eastern and Southern Africa (Figure S1 and Table S1). We focused on seven geographical regions that had more than 20 genomes each, and assigned the remainder to “Other”, including two genomes from Western Africa (Figure 1A). The resulting phylogeny of L2 was divided into two main sublineages: the L2–proto-Beijing and L2–Beijing, supporting previous results (Luo *et al.*, 2015; Shitikov *et al.*, 2017). The L2–proto-Beijing was the most basal L2 sublineage and was restricted to East-and Southeast Asia. L2–Beijing, particularly the “modern” (also known as “typical”) sublineage, was geographically widely distributed and included strains from Africa. We further characterized L2– Beijing using the recently described unified classification scheme for L2 (Shitikov *et al.*, 2017).

**Figure 1.**
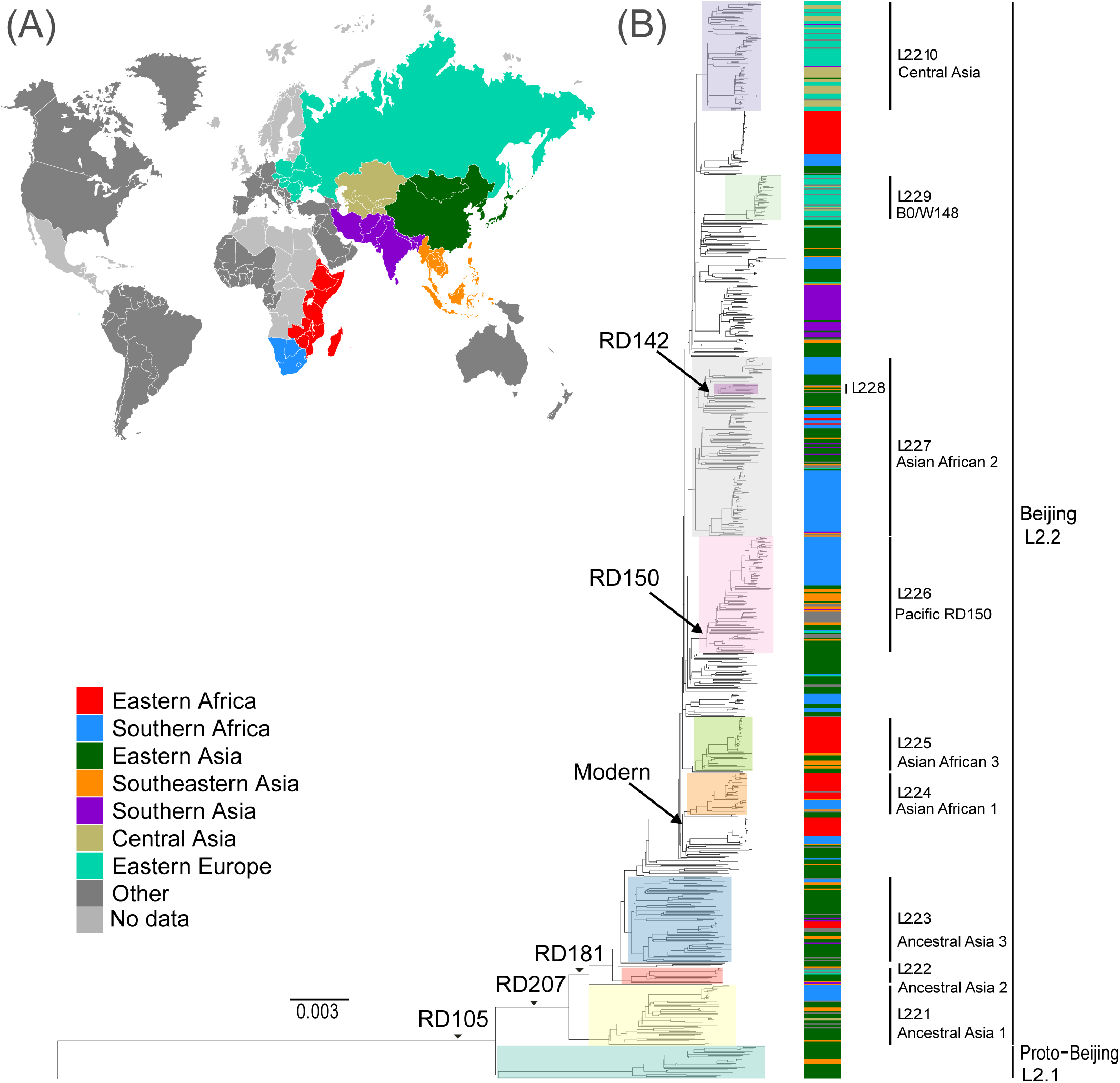
Global phylogeny and geographical distribution of L2 strains. **(A)** Geographical origin (according to United Nations geoscheme) for the 817 L2 strains. The geographical origins with less than 20 strains are colored dark gray and those with missing data light gray. **(B)** Maximum likelihood phylogeny inferred from 33,776 variable single nucleotide positions of the 817 strains. Taxa are colored according to the geographical origin of the strains and the clades are highlighted according to previously defined sublineages. L2 defining markers i.e. deletions (RD) are also mapped onto the phylogeny.

### 2.2 The population structure of L2–Beijing in Eastern and Southern Africa

Our findings showed the population of African L2–Beijing to be heterogeneous (Figure 1B, Figure 2 and Table S2). Most of the African L2–Beijing strains were classified into several groups within the “modern” sublineage, which included primarily the “Asian-African” sublineages (L2.2.4, L2.2.5 and L2.2.7), consistent with previous findings (Merker *et al.*, 2015). We also identified the “ancient” (atypical) strains among the African L2–Beijing. Given that “ancient” L2–Beijing strains (L2.2.1 – L2.2.3) are generally uncommon (Luo *et al.*, 2015), it is interesting to observe such strains in both African regions. In several instances, African L2–Beijing strains did not fall into any of the previously defined groups (Figure 2). Of the two African regions studied here, East Africa had higher proportion of previously uncharacterized L2–Beijing strains (50/101, 50%).

**Figure 2.**
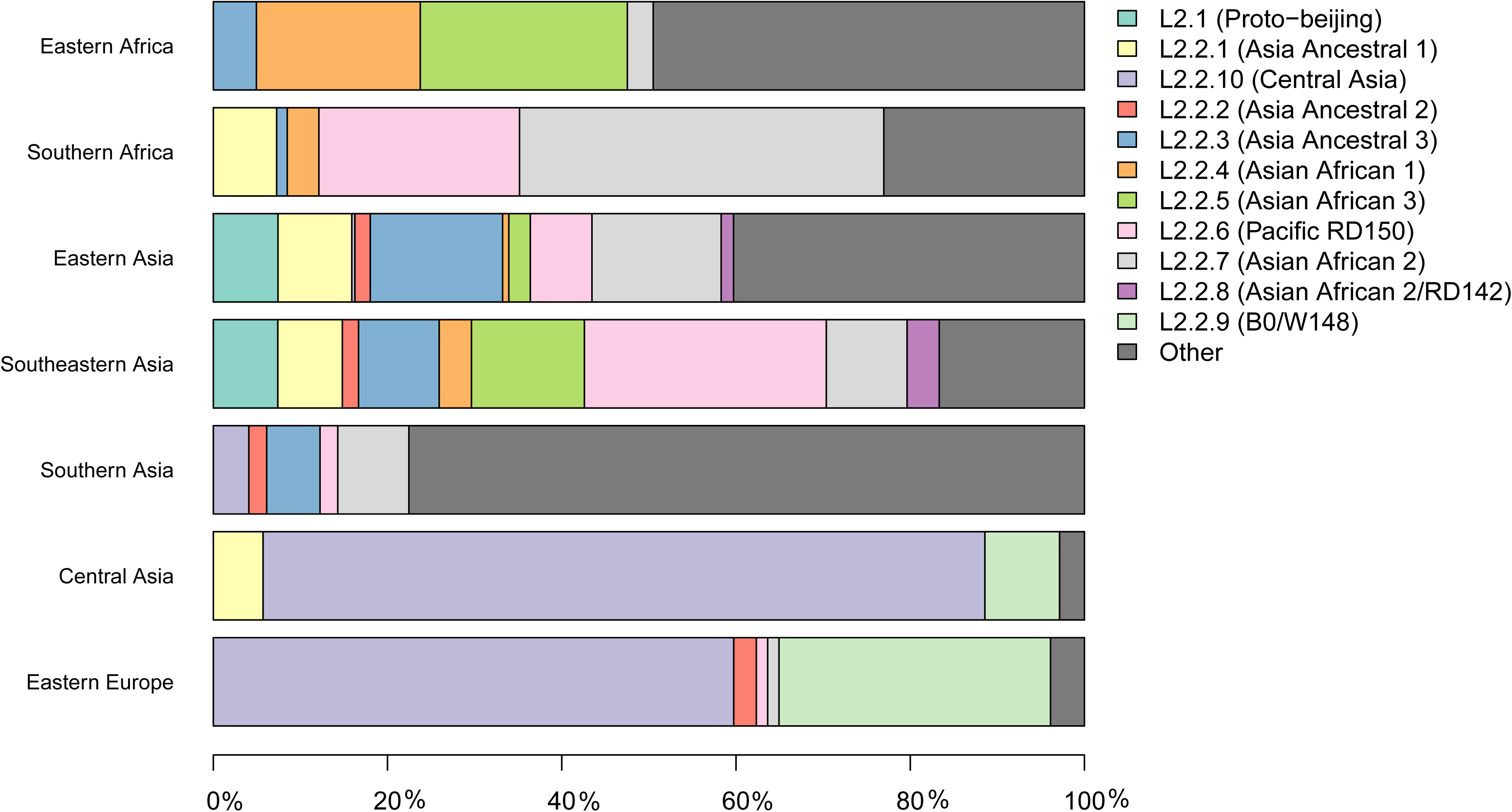
Population structure of L2 strains. Frequency in proportions of L2 sub-lineages across seven geographical regions.

In summary, our findings show that African regions harbored distinct L2–Beijing populations. This is unlike Eastern Europe and Central Asia, where L2–Beijing is dominated by a few highly similar strains (Casali *et al.*, 2014; Eldholm *et al.*, 2016). Of note, L2–Beijing strains typical of Central Asia and Eastern Europe were completely absent from the African populations (Figure 2).

### 2.3 Genetic diversity of L2–Beijing strains across geographic regions

The spatial distribution of L2–Beijing sublineages and the prevalence of “ancient” L2–Beijing strains observed in this study and previously (Luo *et al.*, 2015; Merker *et al.*, 2015), suggest that L2–Beijing has expanded worldwide from Asia. This view can further be supported by the measures of genetic diversity of L2–Beijing in the different geographical regions (Figure 3). As expected, East-and Southeast Asia contained the most genetically diverse L2 populations, which is consistent with previous results (Luo *et al.*, 2015). Conversely, L2 populations in other geographies were less genetically diverse, suggesting recent dissemination of L2 to these regions. Within Africa, Southern Africa showed a higher diversity in L2–Beijing populations compared to Eastern Africa.

**Figure 3.**
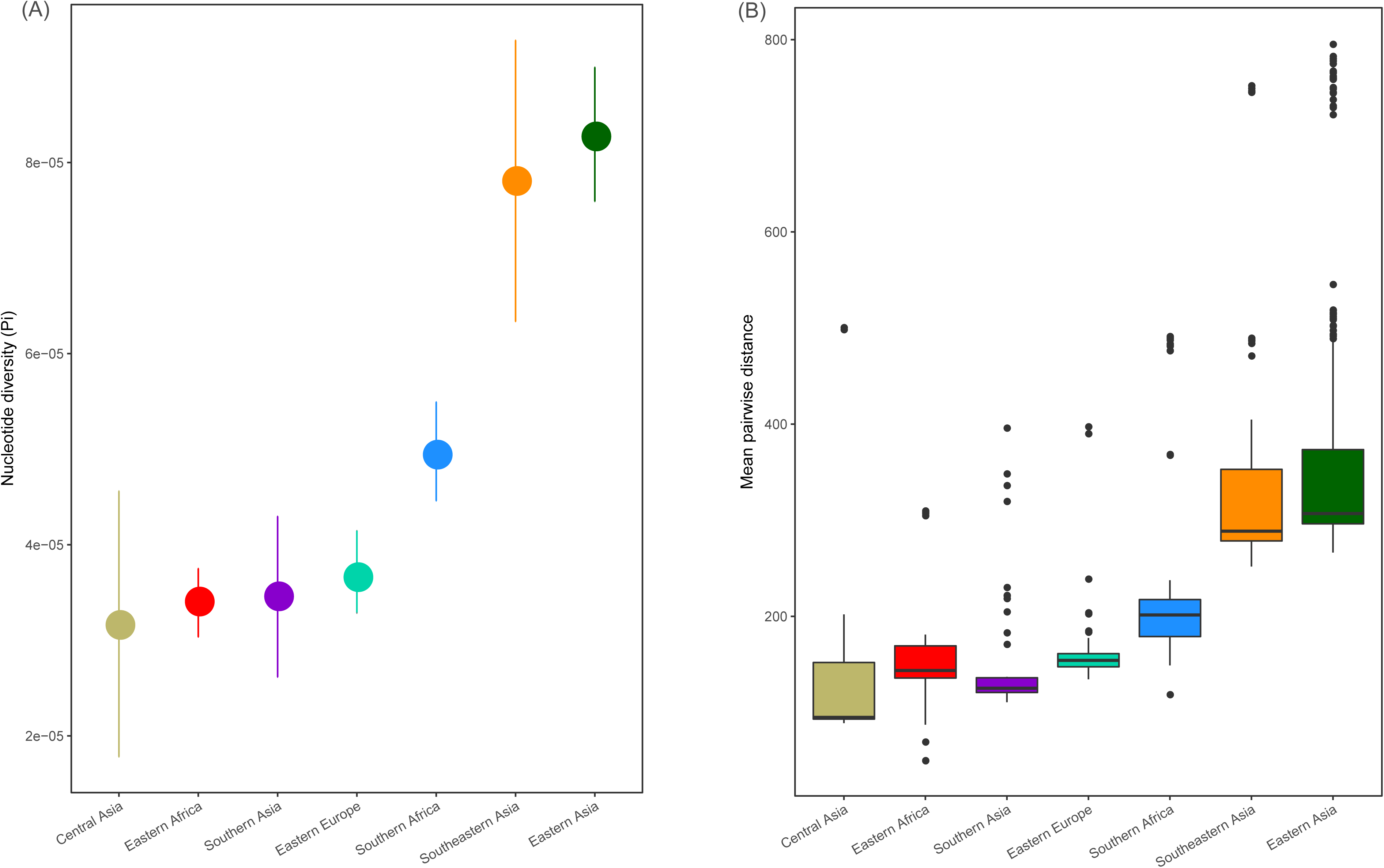
Genetic diversity of L2 strains within geographical regions. **(A)** Nucleotide diversity (π) per site of L2 strains by geography. Error bars are the 95% confidence intervals. **(B)** Pairwise genetic SNP distance of L2 by geography (p values were obtained from Wilcoxon rank-sum tests). Each box represents the 25% and 75% quartiles and the line denotes the median.

The genetic diversity within the African L2–Beijing populations not only reflects the number and variety of source populations but also local patterns of diversification that occurred after their introduction. Therefore, the higher genetic diversity of the L2–Beijing populations in Southern Africa compared to Eastern Africa likely reflects both aspects.

### 2.4 Multiple introductions of L2–Beijing from Asia into Africa

Based on our reconstructed phylogeny, African L2–Beijing strains clustered into several unrelated clades indicating multiple introductions into Africa (Figure 1B). We next investigated the most likely geographical origins of those introductions. As anticipated, our ancestral reconstruction estimated East Asia as the most likely origin of all L2 (posterior probability of 96.7%) and L2–Beijing (posterior probability 92.5%). Our data further indicate that L2–Beijing was introduced into Africa from East-and Southeast Asia on multiple occasions independently. Furthermore, we observed both direct introductions from Asia into Africa as well as subsequent dispersal within the continent (Figure S2 and S3).

Using stochastic mapping, we estimated a total of 13 introductions or migration events (M1 – M13) into Africa (Figure 4). Eight of the African L2–Beijing introductions originated from East Asia and five from Southeast Asia. Out of the 13 introductions, three (M3, M10 and M13) were present in both African regions analyzed here, suggesting initial introductions from Asia followed by subsequent spread within Africa. Overall, our analysis inferred more independent introductions into Southern Africa than Eastern Africa, seven (M1, M4, M7-9, M11 and M12) and three (M2, M5 and M6), respectively. Taken together, our data suggest that multiple migration events have shaped the populations of L2–Beijing in Africa.

**Figure 4.**
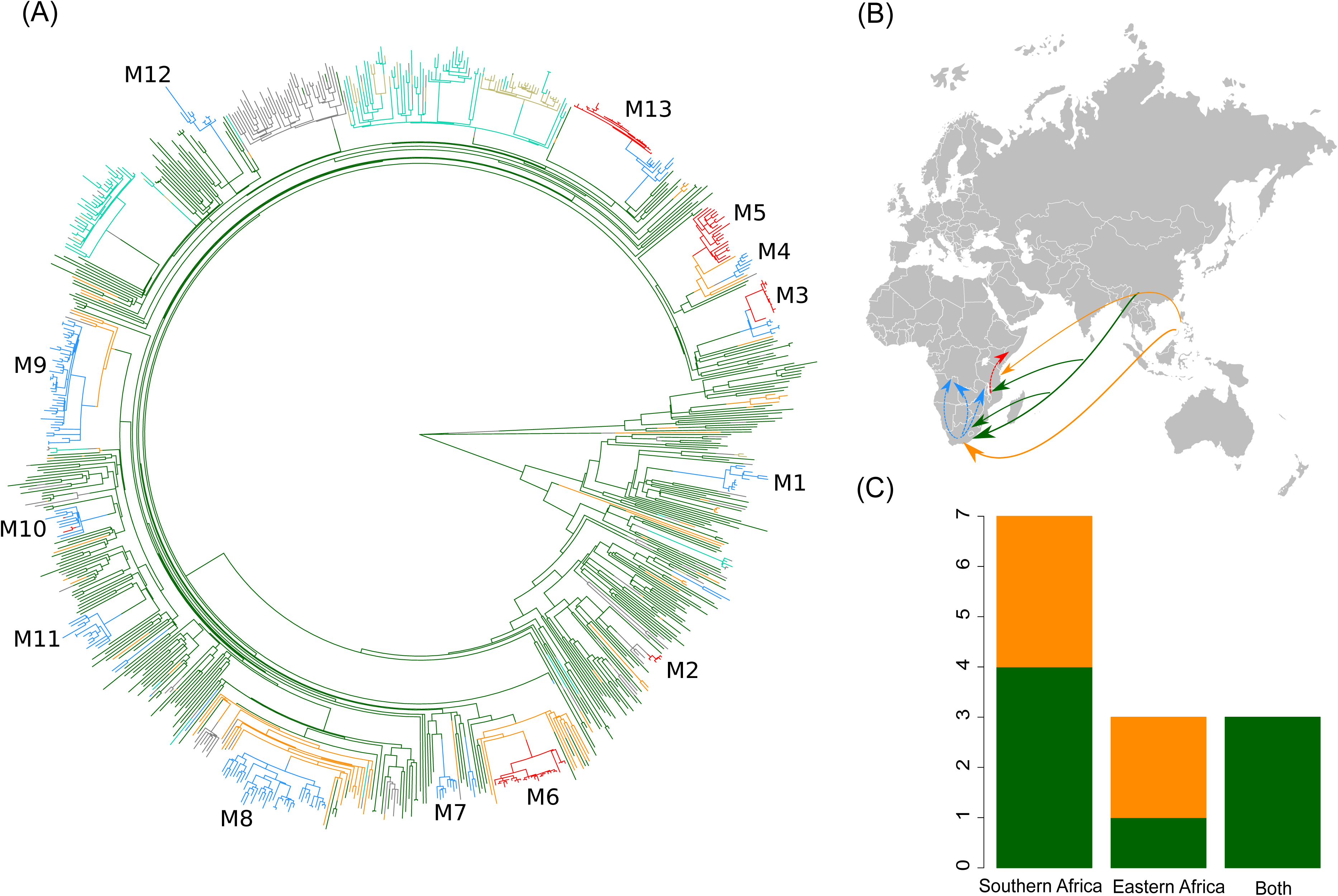
Stochastic mapping of the geographic origin on L2 phylogeny. **(A)** Maximum likelihood phylogeny of the 817 MTB Lineage 2 strains. Branches are colored according to their geographical region inferred from stochastic mapping of geographic origin of L2 strains onto the phylogeny. The 13 migration events to Africa (M1–M13) are indicated. (**B**) Proposed scenario for the multiple introduction events of L2–Beijing into Africa. **(C)** Plot summarizing the number introduction events to Eastern and Southern Africa from East-and Southeast Asia.

### 2.5 Bayesian molecular dating

Different hypotheses have been formulated on the possible timing of the introduction of L2–Beijing into Africa (van Helden *et al.*, 2002). Here we used tip-calibration to date the phylogenetic tree of L2 and estimate the age of its introduction to Africa. For these analyses, we identified 308 strains among the 817 for which the sampling year was known. These strains were sampled during a period of 19 years; 1995 - 2014 (Figure S4), were evenly distributed on the complete phylogenetic tree (Figure S5) and included 40% members of the African L2–Beijing strains (Figure S6). Eleven of the 13 African introductions were represented in this dataset (M1-M3 and M6-M13).

We detected no overlap in the 95% credibility interval of the clock rate estimates of observed and randomized datasets indicating that there was sufficient temporal signal in the dataset to perform inference (see methods, Figure S7). Further, We found that the UCLD clock had the highest marginal likelihood and a Bayes Factor of 27 with the second best fitting model, the strict clock (Table 1), indicating strong evidence in favor of the UCLD clock (Kass and Raftery, 1995).

**Table 1:** Model selection based on path sampling Log-Marginal Likelihood

| Clock model | Log(e) Marginal Likelihood | Bayes factor (UCLD vs model) |
| --- | --- | --- |
| **UCLD** | -5374827 |  |
| **Strict** | -5374854 | 27 |
| **UCED** | -5374897 | 70 |

We performed a phylogenetic analysis with BEAST2 using the UCLD clock, and repeated the tip randomization test (see methods). We found that the 95% credibility interval of the clock rate estimates of observed and randomized datasets did not overlap, confirming that there was a temporal signal in our sequence data (Figure S8). Additionally, under the UCLD model, the coefficient of variation (COV), which is a summary of the branch rates distribution (standard deviation divided by the mean), gives an indication on the clock-likeness of the data (Drummond *et al.*, 2006). A coefficient of variation of zero indicates that the data fit a strict clock, whilst a greater COV indicates a higher heterogeneity of rates through the phylogeny. We obtained a mean COV of 0.22 (95% credibility interval= 0.1732, 0.2732), indicating a moderate level of rate variation across different branches and thus supporting the results of the path sampling analysis that favored the UCLD model.

### 2.6 Recent origins of the African L2–Beijing clades

We used the UCLD clock model to infer the clock rate and divergent times of the 308 L2 strains with known sampling dates and estimated a mean substitution rate of 1.34 × 10^−7^ [95% Highest Posterior Density (HPD), 9.2867 × 10^−^8 - 1.7719 × 10^−7^]. These estimates are in agreement with previously reported rates from epidemiological studies (Walker *et al.*, 2013; Eldholm *et al.*, 2015). We estimated the most recent common ancestor (MRCA) of the extant L2–Beijing of the 308 strains to the year 1225 [95% HPD, 900 - 1519] (Figure S9). For each African clade, we estimated the year of introduction using the 0.975 quantile of the HPD of the age of the MRCA as the upper limit (most recent possible year) and the 0.025 quantile of the HPD of the divergence time between the closest non-African L2–Beijing strain (the closest outgroup) and the African clade of interest as lowest limit (most ancient possible year). Our estimates placed the earliest introductions of the African L2– Beijing (M1, M3, M7, and M12) in the 18^th^ and 19^th^ century (Figure 5 and Table S3). Four additional migration events (M6, M9, M10, and M11) were estimated to have occurred between the beginning of the 19th century and the first half of the 20^th^ century. Finally, the three most recent introductions to Africa happened in the second half of the 20^th^ century (M2, M8, and M13). Diversity patterns of the African clades exclusive to Eastern-and Southern Africa could further provide support for the recent introductions of African L2–Beijing. We thus calculated the pairwise SNP distances within the individual introductions to explore the local patterns of diversification associated with regional epidemics after the introductions. Although strains within Southern African introductions were relatively more distantly related, L2–Beijing strains from both African regions were on average 20 to 40 SNPs apart (Figure S10 and S11). The latter thresholds were proposed to correspond to strains involved in transmission clusters of estimated 50 to100 years (Meehan *et al.*, 2018), supporting the relatively recent introductions of L2–Beijing into the African continent.

**Figure 5.**
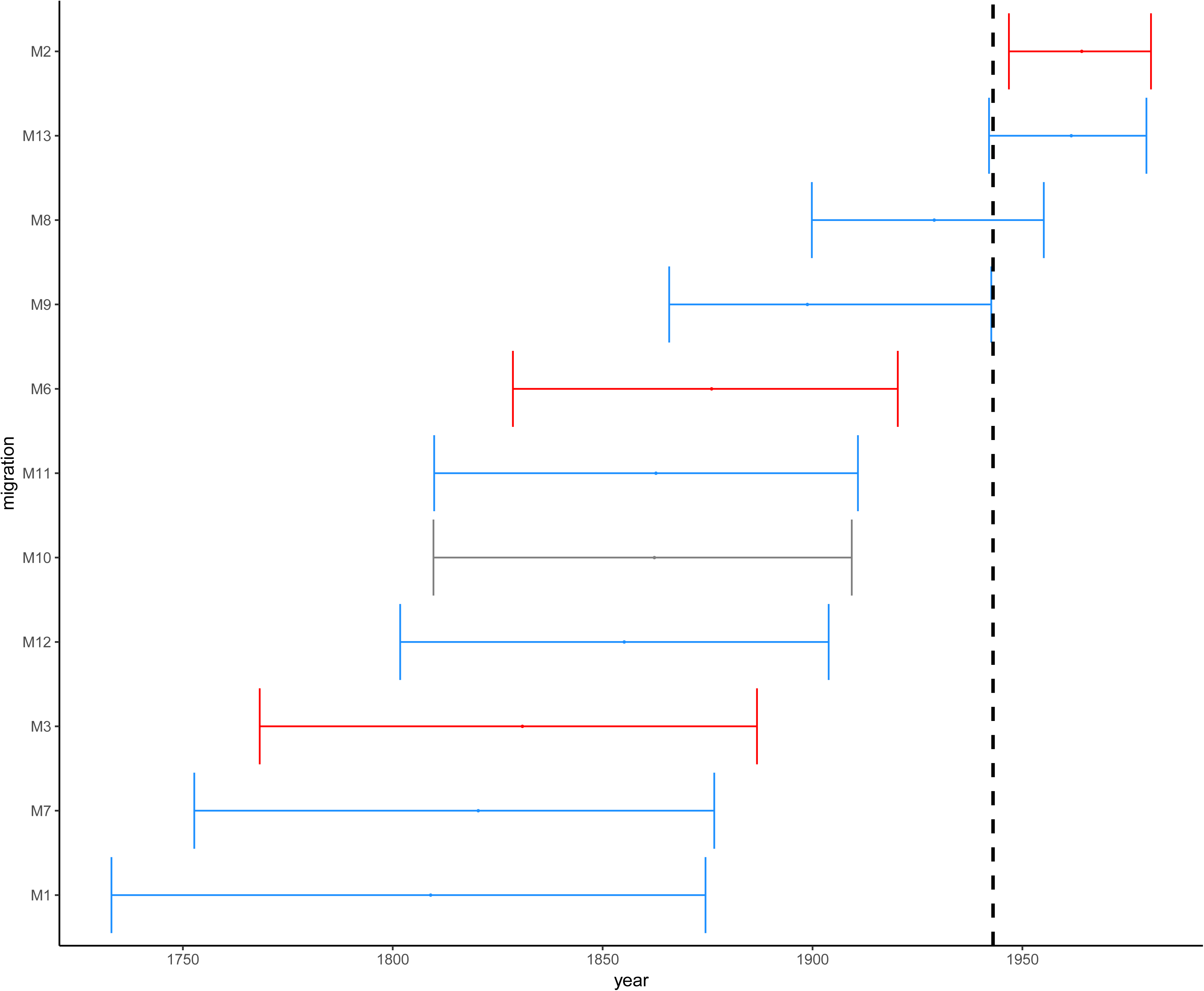
Estimated time in median ages for the introductions of African L2–Beijing (M1-M3 and M6-M13). Introductions to Eastern Africa are colored in red and those to Southern Africa in blue. Migration M10 contained L2–Beijing from both Southern and Eastern Africa. Dotted line marks the year of first anti-TB drug discovery (1943). The error bars correspond to the 95% HPD.

Overall, these results indicate that the different introductions of L2–Beijing to Africa occurred over a period of 300 years. While the earliest introduction is unlikely to have happened after 1732 - 1874, the most recent is unlikely to have occurred before 1946 - 1980.

### 2.7 Introductions of L2–Beijing into Africa unrelated to drug resistance

Because of the repeated association of L2–Beijing with antibiotic resistance (Borrell and Gagneux, 2009), the emergence and dissemination of L2–Beijing strains has been attributed to drug resistance. However, our estimated timing of these introductions suggest that African L2–Beijing strains were introduced prior the discovery of TB antibiotics, and thus must have involved drug-susceptible strains (Figure 5). To explore this question further, we assessed the drug resistance profiles of L2– Beijing strains linked to the various introduction events into the two African regions. We found that all the Eastern African populations contained only drug-susceptible strains and that approximately three-quarters of L2–Beijing strains in the Southern African populations were drug-susceptible, with the remaining being either mono–, multi– or extensively drug-resistant (Figure 6 and Figure S12). Taken together, these results suggest that the emergence of L2–Beijing in Africa, particularly in Eastern Africa, was not driven by drug resistance. Moreover, our data indicate independent acquisition of drug resistance for the resistant strains detected in the Southern African L2–Beijing population (Figure 6), which might partly contribute to the subsequent spread of L2–Beijing in Southern Africa but not in Eastern Africa.

**Figure 6.**
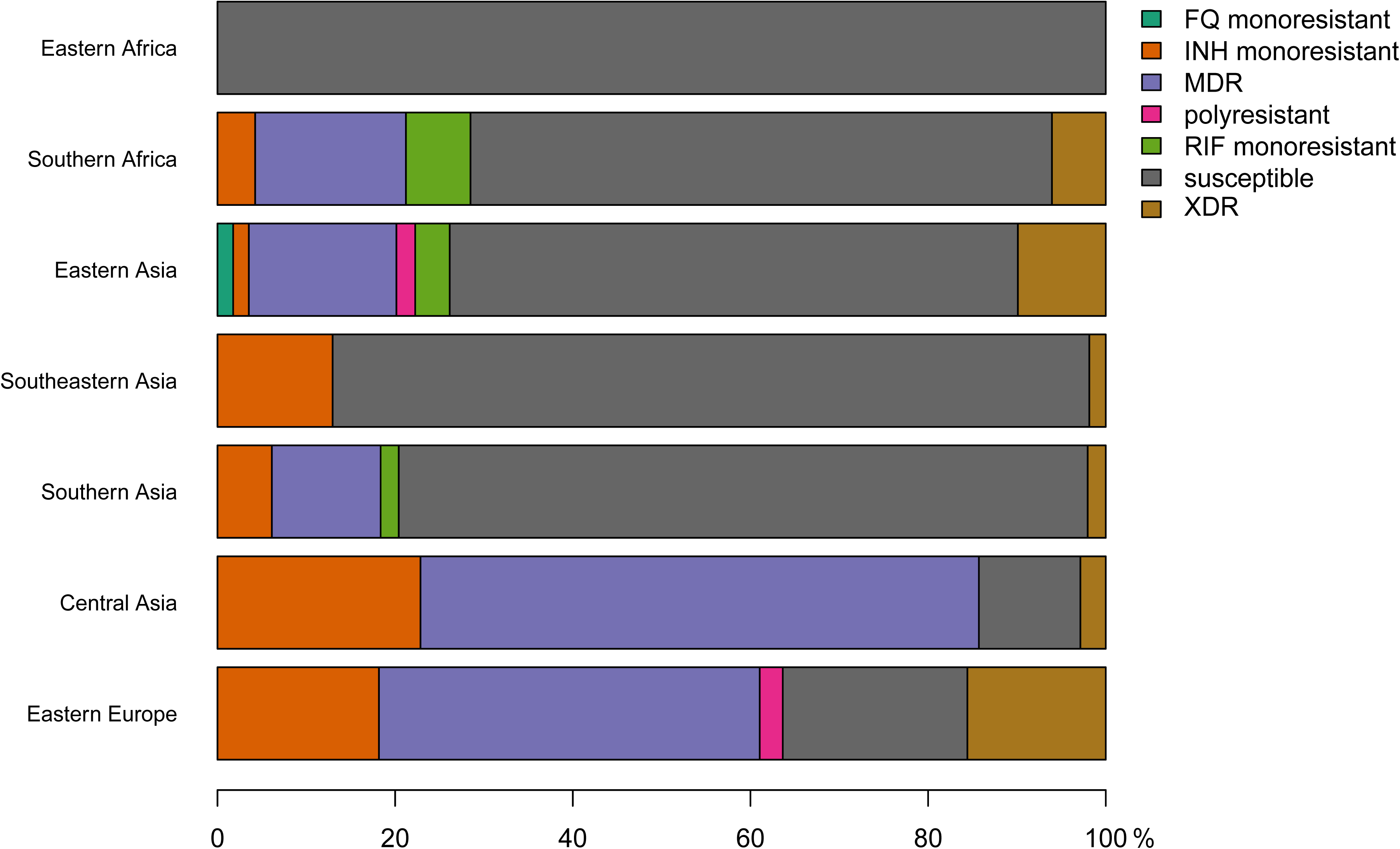
Proportions of drug resistance profiles for L2 strains in seven geographical regions.

## 3 Discussion

This study investigated the most likely geographical origin of the L2–Beijing in Africa. In line with previous findings (Luo *et al.*, 2015; Merker *et al.*, 2015), we identified East Asia as the most likely place of origin of L2 and L2–Beijing. Our findings further revealed multiple independent introductions of L2–Beijing into Africa linked to separate populations originating from both East-and Southeast Asia. Some of these introductions were followed by further onward spread of L2– Beijing within African regions. Finally, we demonstrate that most introductions of L2–Beijing on the continent occurred before the age of antibiotics.

L2–Beijing has received much attention given its hypervirulence in infection models (Manca *et al.*, 2001; Ribeiro *et al.*, 2014), faster progression to disease and higher transmission potential in humans (de Jong *et al.*, 2008; Holt *et al.*, 2018), frequent association with drug resistance, and recent emergence in different regions of the world (Bifani *et al.*, 2002; Borrell and Gagneux, 2009; Fenner *et al.*, 2013). Several studies indicate L2–Beijing originated in Asia and spread from there to the rest of the world (Luo *et al.*, 2015; Merker *et al.*, 2015). Our results support this notion by identifying “Asia” as the most likely geographical origin of both L2 and L2–Beijing based on our ancestral reconstructions and the fact the L2–Beijing populations in Asia are much more diverse than in other regions. In addition, our findings show that L2–Beijing was introduced into Africa multiple times from both East-and Southeast Asia. The presence of L2–Beijing in South Africa has previously been proposed to be due to the importation of slaves from Southeastern Asia by Europeans in the 17^th^ and 18^th^ centuries followed by the import of Chinese labor-forces in the early 1900s (van Helden *et al.*, 2002; Mokrousov *et al.*, 2005). Our Bayesian dating estimates predicted the earliest introductions of L2– Beijing into Africa to have occurred in the 18^th^ and 19^th^ centuries, concurring with these proposed time periods. However, our findings also point to later introductions of L2–Beijing into the continent in the 19^th^ and early 20^th^ centuries. The timings of the latest three introductions in the second half of the 20^th^ century coincide with the decolonization and post-colonial period in Africa when investments into infrastructure and other projects by Chinese enterprises substantially increased (Yuan, 2006; Rice, 2011). These activities also brought many Chinese workers to Africa during a time when TB was still very prevalent in China (Murray, 2018). Hence, many of these workers were likely latently infected with L2–Beijing and might have later reactivated (Moreira Pescarini *et al.*, 2017). Overall, our findings suggest that L2–Beijing has emerged in Africa over the last 300 years and that these introductions have occurred sporadically ever since.

The repeated association of L2–Beijing with drug resistance (Borrell and Gagneux, 2009) has led some to propose that drug resistance is another reason why this sublineage might successfully compete against and eventually replace other *M. tuberculosis* genotypes (Parwati, Van Crevel and Van Soolingen, 2010). However, the underlying reasons for the association of L2–Beijing with drug resistance remains unclear (Borrell and Trauner, 2017), and it is also far from universal, with several reports from e.g. China and other regions finding no such association (Hanekom *et al.*, 2007; Yang *et al.*, 2012). Our results show that most introduction events of L2–Beijing into Africa pre-date the antibiotic era, and because of that these introductions were most likely caused by drug-susceptible strains. The notion that the initial emergence of L2–Beijing in Africa was not driven by drug resistance is further supported by our findings that none of L2–Beijing strains from Eastern Africa strains analyzed here were drug-resistant. Of note, our observations suggest that drug resistance in South Africa was acquired via independent events post initial introductions from Asia. This is in sharp contrast to the situation in Eastern Europe and Central Asia, where L2–Beijing is highly prevalent but dominated by few recently expanded drug-resistant clones, which account for up to 60% of the L2–Beijing populations in some of these countries (Casali *et al.*, 2014; Eldholm *et al.*, 2016). The association of L2–Beijing with drug resistance in these regions were likely favored by the economic and public health crises that followed the collapse of Soviet Union (Luo *et al.*, 2015; Merker *et al.*, 2015).

Based on our finding that the original introductions of L2–Beijing into Africa involved drug-susceptible strains and that the prevalence of drug-resistant L2–Beijing in Africa overall is comparably low (WHO, 2017), we propose that some of the other characteristics of this sub-lineage, in particular its high virulence, high transmissibility and rapid progression from infection to disease, were responsible for the initial competitive success of L2–Beijing in Africa. Given that the MTBC overall originated in Africa (Wirth *et al.*, 2008; Comas *et al.*, 2013), TB epidemics on the continent were caused by many different “native” genotypes prior to foreign contacts (Comas *et al.*, 2015). The emergence and expansion of “foreign” genotypes including L2–Beijing post-contact demonstrate their ability to successfully compete against the existing genotypes on the continent, irrespective of drug resistance. Following their initial establishment, poor TB treatment programs subsequently selected for drug resistance in L2–Beijing but also in other MTBC lineages, which might have facilitated their further spread in countries such as South Africa (Müller *et al.*, 2013).

This study is limited by the fact that we analyzed a globally diverse collection of L2 genomes available in public repositories. Hence, these strains might not be fully representative of the respective geographical regions. Moreover, our African L2–Beijing dataset came from convenient sampling and comprised L2–Beijing mainly from Eastern and Southern Africa, as whole genome data of L2–Beijing from the other African regions were unavailable at the time of the study. However, the representation of African L2–Beijing in our sample reflects the overall prevalence of this sub-lineage as recently reported for the continent (Mbugi *et al.*, 2016; Chihota *et al.*, 2018). Moreover, although regions outside of Eastern-and Southern Africa were underrepresented, this is unlikely to invalidate our findings regarding the multiple independents of L2–Beijing into Africa, except by underestimating the number of true introductions.

In conclusion, this is the first study to address the geographical origins of L2–Beijing in Africa using whole genome sequencing data. Our findings indicate multiple independent introductions of L2– Beijing epidemics into Africa from East-and Southeast Asia during the last 300 years that were unrelated to drug resistance. The TB epidemics in Africa have remained fairly stable over the last few decades (WHO, 2017). However, Africa’s population growth and increasing urbanization (driven by booming economies) are likely to have an impact on the future of TB in this continent, whether directly by e.g. facilitating transmission or indirectly by promoting new risk factors such as diabetes that increase TB susceptibility (Dye and Williams, 2010). It is therefore crucial to follow the TB epidemics in the continent very closely, especially those related to hypervirulent strains such as L2– Beijing, as these might take particular advantage of this expanding ecological niche (Cowley *et al.*, 2008).

## 4 Material and Methods

### 4.1 Identification of Lineage 2 strains and whole-genome sequencing

We obtained whole-genome sequencing data of L2 strains from the two previously largest studies focusing on the evolutionary history and global spread of L2–Beijing strains (Luo *et al.*, 2015; Merker *et al.*, 2015). We then identified additional published genomes as African representatives of L2–Beijing strains from other studies (Guerra-Assunção *et al.*, 2015; Manson *et al.*, 2017). Moreover, we newly sequenced 116 additional L2–Beijing strains using Illumina HiSeq 2000/2500 paired end technology (PRJNA488343). In total, we included 817 L2 genome sequences (Figure S1 and Table S1).

### 4.2 Whole genome sequence analysis and phylogenetic inference

We used a customized pipeline previously described to map short sequencing reads with BWA 0.6.2 to a reconstructed hypothetical MTBC ancestor used as reference (Comas *et al.*, 2013). SAMtools 0.1.19 was used to call single nucleotide polymorphisms (SNPs), and these SNPs were annotated using ANNOVAR and customized scripts based on the *M. tuberculosis* H37Rv reference annotation (AL123456.2). For downstream analyses, we excluded SNPs in repetitive regions, those annotated in problematic regions such as ‘PE/PPE/PGRS’ and SNPs in drug-resistance associated genes. Small insertions and deletions were also excluded from the analyses. Only SNPs with minimum coverage of 20x and minimum mapping quality of 30 were kept. All SPNs classified by Samtools as having frequencies of the major non-reference allele lower than 100% (AF1<1) within each genome were considered to be heterogeneous and were treated as ambiguities, and otherwise considered fixed (AF1=1). We concatenated fixed SNPs from the variable positions obtained, which yielded a 33,776 bp alignment. The alignment was then used to infer a maximum likelihood phylogeny using RaxML 8.3.2 with a general time reversible (GTR) model in RAxML and 1,000 rapid bootstrap inferences, followed by a thorough maximum-likelihood search (Stamatakis, 2006).

### 4.3 Phylogeographic analyses

#### 4.3.1 Reconstruction of ancestral state

To investigate the likely geographic origin of L2–Beijing strains in Africa, we inferred the historical biogeography of L2 using the RASP software (Yu *et al.*, 2015) on a representative subset of 430 genomes due to software’s sample limitation. We achieved this subset by performing hierarchical clustering implemented in *pvclust* package in R (Suzuki and Shimodaira, 2006) on a distance matrix and randomly removing clustered genomes. We applied a Bayesian based method in RASP to reconstruct ancestral geographical states on the phylogeny of 430 L2 genomes. We used geographical regions as proxy for origins of the L2 strains and loaded them as distributions. We then ran Bayesian analysis with 5 chains and 500 generations.

#### 4.3.2 Stochastic character mapping

To determine the number of introduction events of L2–Beijing into African regions, we applied stochastic mapping (SIMMAP) on the 817 L2 phylogeny using phytools package 0.6.44 in R (Revell, 2012). Geographical origin of the L2 strains was treated as a discrete trait and modeled onto the phylogeny using ARD model with 100 replicates. This model permits independent region-to-region transfer. We referred to the resulting introductions as migration events “M”, considering only those introductions with more than 5 genomes.

#### 4.3.3 Population genetic analyses

##### 4.3.3.1 Nucleotide diversity (pi)

We calculated the mean pair-wise nucleotide diversity per site (Pi) measured by geographic region. We excluded geographic regions represented by less than 20 genomes. Confidence intervals were obtained by bootstrapping through resampling using the sample function in R with replacement and the respective lower and upper confidence levels by calculating 2.5th and 97.5th quartiles.

##### 4.3.3.2 Pairwise SNP distances

We used *dist.dna* function of ape package implemented in R (Paradis, Claude and Strimmer, 2004) to calculate pairwise SNP distances with raw mutation counts and pairwise deletions for gaps. Mean pairwise SNP distance to all strains of the same geographic population was calculated per strain and the distribution of the mean SNP pairwise distance plotted. The mean pairwise SNP distances were assumed not to be normally distributed and we therefore used Wilcoxon rank-sum test to test the differences among geographic regions. Additionally, we calculated pairwise SNP distances within African L2–Beijing populations for migration events with more than 10 genomes each.

#### 4.3.4 Drug Resistance

To distinguish between drug-susceptible and drug-resistant strains, we used genotypic drug resistance molecular markers previously described (Steiner *et al.*, 2014). We categorized strains into: susceptible as having no drug resistance specific mutations; monoresistant as having mutations conferring resistance to a single drug; MDR as having mutations conferring resistance to isoniazid and rifampicin; and extensively drug-resistant (XDR) as having mutations conferring resistance to fluoroquinolones and aminoglycosides in addition to being MDR (Table S4).

#### 4.3.5 Bayesian molecular dating

##### 4.3.5.1 Data preparation and preliminary analysis

To estimate the historical period in which L2–Beijing was introduced to Africa, we performed a set of Bayesian phylogenetic analyses using tip-calibration (Rieux and Khatchikian, 2017). Among the 817 studied L2 strains, we had information on the year of sampling for 308. We performed all further analysis on this subset of 308 strains. We excluded all genomic positions that were invariable in this subset and all positions that were undetermined (missing data or deletions) in more than 25% of the strains, and obtained an alignment of 10,769 polymorphic positions.

In tip dating analysis it is important to test whether the dataset contains strong enough temporal signal (Rieux and Balloux, 2016). To do this, we performed a tip randomization test (Ramsden *et al.*, 2008) as follows. We used BEAST2 v. 2.4.8 (Bouckaert *et al.*, 2014) to run a phylogenetic analysis with a HKY + GAMMA model (Hasegawa, Kishino and Yano, 1985), a constant population size prior on the tree and a strict molecular clock. Additionally, we used the years in which the strains were sampled to time-calibrate the tree, and we modified the extensible markup language (xml) file to specify the number of invariant sites as indicated by the developers of BEAST2 here: https://groups.google.com/forum/#!topic/beast-users/QfBHMOqImFE (*strict_preliminary.xml*). We ran three independent runs (245 million generations in total), and we used Tracer 1.7 (Rambaut *et al.*, 2018) to identify the burn-in (8 million generations), to assess that the different runs converged, and to estimate the effective sample size (ESS) for all parameters, the posterior and the likelihood (ESS > 110 for all parameters). We then used TipDatingBeast (Rieux and Khatchikian, 2017) to generate 20 replicates of the xml file in which the sampling dates were randomly reassigned to different strains. For each replication, we ran the same BEAST2 analysis as for the original (observed) dataset (one run per replicate, 50 million generations, 10% burn-in). We used TipDatingBeast to parse the log files output of BEAST2 and compare the clock rate estimates for the observed data and the randomized replications. The estimates of the molecular clock rate did not overlap between the observed and the randomized dataset, indicating that there is a clear temporal signal and that we could proceed with further analysis (Figure S5).

##### 4.3.5.2 Model selection

To identify the model that best fits the data, we estimated the marginal likelihood of three different clock models: UCED and UCLD (Drummond *et al.*, 2006). We used the Model selection package of BEAST2 to run a path sampling analysis (Lartillot and Philippe, 2006) following the recommendations of the BEAST2 developers (http://www.beast2.org/path-sampling/). We used the following settings: 100 steps, 4 million generations per step, alpha = 0.3, pre-burn-in = 1 million generations, burn-in for each step= 40% (*\*PS.xml*). For these analyses, we used proper priors as suggested by (Baele *et al.*, 2012).

##### 4.3.5.3 UCLD analysis

Since the model selection analysis indicated that the UCLD clock was the best fitting model, we repeated the analysis using the UCLD and the same settings used in the path sampling analysis, sampling every 10,000 generations. We ran three independent runs (800 million generations in total), we used Tracer 1.7 (Rambaut *et al.*, 2018) to identify the burn-in (10 million generations), to assess that the different runs converged and to estimate the effective sample size (ESS) for all parameters, the posterior and the likelihood (ESS > 260 for all parameters) (*UCLD_final.xml* and Table S5)

We checked the sensitivity to the priors by running one analysis of 250 million generation sampling from the prior, and compared the parameter estimates with the analysis using the data. We observed the posterior distribution and the prior distribution of all parameters are very distinct (Table S6), indicating that the parameter estimates are influenced by the data and not by the priors (Bromham *et al.*, 2018).

We repeated the tip randomization test with the UCLD model as described above (20 replicates, one run per replicate, 105 million generations per replicate or more, burn-in 10%), and again we found a temporal signal (Figure S8).

To summarize the results, we sampled the trees from the three runs (5% burn-in corresponding to 10 million generations or more, sampling every 25,000 generation). We then summarized the 31,758 sampled trees, created a maximum clade credibility tree using the software TreeAnnotator from the BEAST2 package and used FigTree version 1.4.2 for visualization (Figure S9).

## 5 Conflict of Interest

We declare no conflict of interest.

## 6 Author Contributions

LKR, DB, FM, DS, LF and SG planned the study, SDL, BM and JF performed the experiments, LKR, DB, FM, SMG, SDL, BM, CB, SB, KM, MB, LJ, KR and LF contributed strains and prepared the data, LKR, DB, FM and SG analyzed the data, LKR, DB, FM and SG drafted the manuscript. All authors critically reviewed the manuscript.

## 7 Acknowledgments and funding information

We would like to thank Sebastián Duchêne and Yan Yu for their technical support and Linda-Gail Bekker for contributing strains. All bioinformatics analyses were performed at the scientific computing core facility of the University of Basel, sciCORE (http://scicore.unibas.ch/). This work was supported by the Swiss National Science Foundation (grants 310030_166687 to SG), the European Research Council (309540-EVODRTB to SG) and SystemsX.ch This research was also partially supported (strain collection) by a funding supplement from the National Institutes of Allergy and Infectious Diseases (NIAID) under award numbers U01 AI069924 (IeDEA Southern Africa) and U01 AI069911 (IeDEA East Africa).

## Data Availability Statement

The additional datasets i.e. additional xml files and scripts for this study can be found here https://github.com/SwissTPH/TBRU_L2Africa.git

